# LysM: a bacterial self-encoded peptidoglycan hydrolase as an antimicrobial and anti-biofilm agent for Gram-negative nosocomial pathogens

**DOI:** 10.64898/2026.10.20.760325

**Authors:** Yu Zhang, Dejian Liu, Min Zhao, Lanxin Wang, Yanhong Huang, Haoyu Liu, Jingchao Zhang, Bo Ma, Di Wang, Kun Zhao, Jintao Liu, Liang Li, Luyan Z. Ma

## Abstract

Agents with both anti-biofilm property and broad-spectrum antimicrobial activity are urgently needed, especially for drug-resistant nosocomial pathogens. Here, we report a bacterial genome-encoded glycoside hydrolase, named LysM, with lytic activity against multiple Gram-negative nosocomial pathogens and little toxicity to human cells. 3D structural modelling and site-directed mutagenesis indicates that LysM is a peptidoglycan-specific glycoside hydrolase. In combination with EDTA, exogenous LysM induces explosive bacterial cell lysis, exhibits potent bactericidal activity, and eliminates 24 h-old biofilms formed by Gram-negative ESKAPE group pathogens with the highest activity on three nosocomial pathogens, namely *Pseudomonas aeruginosa, Acinetobacter baumannii,* and *Enterobacter cloacae,* all of which contain LysM homologs in their genome. More strikingly, LysM supplied in growth media without EDTA, also inhibits the growth of several nosocomial pathogens. Single-cell tracking analysis reveals that LysM completely prevent biofilm formation of *P. aeruginosa* by reducing bacterial surface-adherence and extending the doubling time. A LysM-based “cocktail” consisting of LysM, EDTA, and matrix exopolysaccharide-degrading glycosidase PslG, shows potent anti-biofilm activity against *P. aeruginosa*, including drug-resistant clinical isolates, both *in vitro* and in human airway organoids (HAOs) biofilm infection model. In addition, LysM enhances the efficacy of many antibiotics from all major classes, including colistin, tobramycin, ciprofloxacin, and tetracycline. Our findings not only provide a clinical candidate for the treatment of nosocomial infections but also open a new path for antimicrobial discovery and anti-biofilm strategies, as homologues of LysM are present in the genomes of many bacteria and phages.

## Introduction

Antibiotic resistance in pathogens is one of the most serious global public health threats, particularly among the ESKAPE group, comprising *Enterococcus faecium*, *Staphylococcus aureus*, *Klebsiella pneumoniae*, *Acinetobacter baumannii*, *Pseudomonas aeruginosa*, and *Enterobacter* specie*s* (1). These bacteria commonly cause nosocomial infections and are key priority pathogens listed by the World Health Organization for research & development of new antimicrobials (2–4). The development of multidrug resistance (MDR) compromises antibacterial chemotherapy, particularly for Gram-negative bacterial pathogen-associated infections (2–4). It has been estimated that bacterial MDR was directly responsible for 1.27 million global deaths in 2019 and contributed to 4.95 million deaths (5). The rapid spreading of MDR Gram-negative pathogens leaves clinicians few choices from the antibiotic pipeline (6, 7). Consequently, novel antimicrobials and alternative strategies are desperately needed. In this decade, it has been widely recognized that matrix-enmeshed microbial communities, termed as biofilms, are extreme tolerance to almost all antibiotic classes (8–11). In addition, biofilm bacteria are also protected from the host immune response, giving rise to chronic infections that are notoriously difficult to eradicate and become a source of drug resistance bacteria (12–14). Therefore, antimicrobials with anti-biofilm activity have been a new paradigm for antimicrobial discovery.

Antimicrobials targeting on the bacteria cell wall, especially peptidoglycan (PG) have a low propensity for pathogens to generate spontaneous resistance (15). Given that PG is a unique component of bacterial cell, the PG-targeting antimicrobials are usually harmless to human or animal cells and much less toxic compared to antimicrobials targeting to the cell membrane (16). Bacteriophage endolysin is a type of enzyme that can damage bacterial cell wall through PG hydrolysate, which has been recognized as a new class of antimicrobials. Numerous endolysins have been applied in many fields to combat MDR bacteria over the past decade (16–19). However, it remains largely required for discovery of new endolysins due to the narrow antimicrobial spectrum of endolysin.

*P. aeruginosa* is an opportunistic pathogen that can cause life-threatening infections in patients with cystic fibrosis and individuals with an immune-compromised system (20–23). Infections caused by this pathogen are notoriously difficult to eradicate due to its ability to form biofilms and its antibiotic resistance or tolerance at the individual-cell level. This bacterium can form biofilms on a variety of surfaces such as the mucus plugs of the CF lung, contaminated catheters, and contact lenses (24–26). In addition, *P. aeruginosa* represents one of the most common agents isolated from chronic leg ulcers, often as a biofilm resistant to antimicrobial therapy. Our previous study has revealed that glycosyl hydrolase PslG can sensitize biofilm bacteria of *P. aeruginosa* to antibiotics by disrupting the key biofilm matrix exopolysaccharide Psl (27). Turnbull et al. have previously reported that high level expression of a PA0629-encoded protein leads to explosive cell lysis as well as the production of membrane vesicles within a biofilm of *P. aeruginosa* (28). PA0629 is located within a gene cluster that is required for the production of R-type pyocin, the contractile, phage tail-like bacteriocins of *P. aeruginosa* (29). The PA0629-encoded protein is a homologue of endolysin.

In this study, we investigate whether the PA0629-encoded protein, herein named as LysM, can lyse multiple bacteria when supplied exogenously and whether it possesses anti-biofilm activity. Strikingly, LysM can hydrolyze cell wall of Gram-negative ESKAPE pathogens. In combination with EDTA, LysM causes explosive cell lysis and eliminate a formed biofilm. Without EDTA, LysM does not lyse bacteria, but can prevent biofilm formation by interfering with bacterial growth and surface adherence. Furthermore, using a human airway organoid model, we have found that the LysM “cocktail” have the best anti-biofilm efficacy against *P. aeruginosa* biofilm-related infections and this holds true for MDR clinical isolates of *P. aeruginosa*. Lastly, LysM can enhance the efficacy of antibiotics covering major classes against Gram-negative ESKAPE pathogens. Taken together, we identified a bacterial source glycoside hydrolase that has great potential to be an antimicrobial for combating biofilm-related nosocomial infections. Our finding also opens a new avenue for antimicrobial discovery and the development of anti-biofilm strategies.

## Results

### LysM can lyse multiple Gram-negative bacteria when supplied exogenously in combination with EDTA

High level expression of PA0629 in *P. aeruginosa* leads to explosive bacterial cell lysis (28). We wondered whether PA0629-encoded protein could induce the explosive cell lysis of *P. aeruginosa* when supplied exogenously. Thus PA0629-encoded protein (named as LysM hereafter) was expressed in *E. coli* BL21 and purified (23 kDa molecular weight). We first tested the effect of LysM on the cell morphology of *P. aeruginosa* PAO1 in PBS buffer. The LysM alone did not show any visible change on the bacterial cell shape (Fig. S1A). In contrast, upon addition of 0.5 mM EDTA (a concentration that can perturb cell membrane without causing cell death), few bacteria with regular shape were detected post 30 min treatment (Fig. S1A), suggesting that LysM may lyse bacteria under this condition. We then monitored bacterial cells by time-lapse microscopy during the treatment. The LysM and EDTA treatment led to the rod bacteria switching to round shape within 5 min (Fig. 1A and Fig. S1B), and then the round cells subsequently explosive lysed in the next 5-15 min (Video 1, Fig. 1A and B), while a portion of round cells remained the shape during the 30 min of observation (Fig. 1B and Fig. S1B). These results indicated that LysM induced explosive cell lysis when supplied exogenously in combination with EDTA. Consistently, the OD_600 o_f PAO1 declined in the PBS buffer with LysM and EDTA (Fig. 1C). The OD_600 d_ropped below 0.2 post 1.5 h-treatment (the left panel in Fig. 1C) and approximately 98% of bacterial cells were lysed (the right panel in Fig. 1C). The propidium iodide (PI) was employed to detect the dead bacteria during the treatment. The fluorescence intensity of PI significantly increased at the first hour of the LysM and EDTA treatment (Fig. 1D) and along with the increasing of PI-stained bacterial population (Fig. S1C-D), indicating that the treatment of LysM in combination with EDTA caused the bacterial cell death. In contrast, the treatment of PBS, LysM or EDTA alone did not affect either OD_600 (_Fig. 1C) or the PI permeability of PAO1 (Fig. 1D), indicating that LysM is the critical trigger for bacterial death. We further examined the killing effect of LysM on other bacteria. Four nosocomial bacteria, namely *A. baumannii*, *K. pneumoniae, E. coli*, and *Enterobacter cloacae* were examined with the LysM treatment of 2 h (Fig. 1E). Strikingly, *A. baumannii* was highly sensitive to LysM. Post 2 h-treatment, LysM combining with EDTA can kill almost 100% cells of *A. baumannii* and the killing rates of the other three bacteria are all over 75% (Fig. 1E). Surprisingly, LysM alone showed some killing effect on cells of both *A. baumannii* and *E. cloacae* with over 50% of killing rate on *A. baumannii* (Fig. 1E). In addition, *P. putida,* another bacterium of the genus *Pseudomonas*, was not as sensitive as that of *P*. *aeruginosa* to the LysM treatment (Fig. 1E). This PA0629-encoded protein can <u>lys</u>e <u>m</u>ultiple Gram-negative bacteria in the presence of EDTA, thus we named it LysM.

**Fig. 1.**
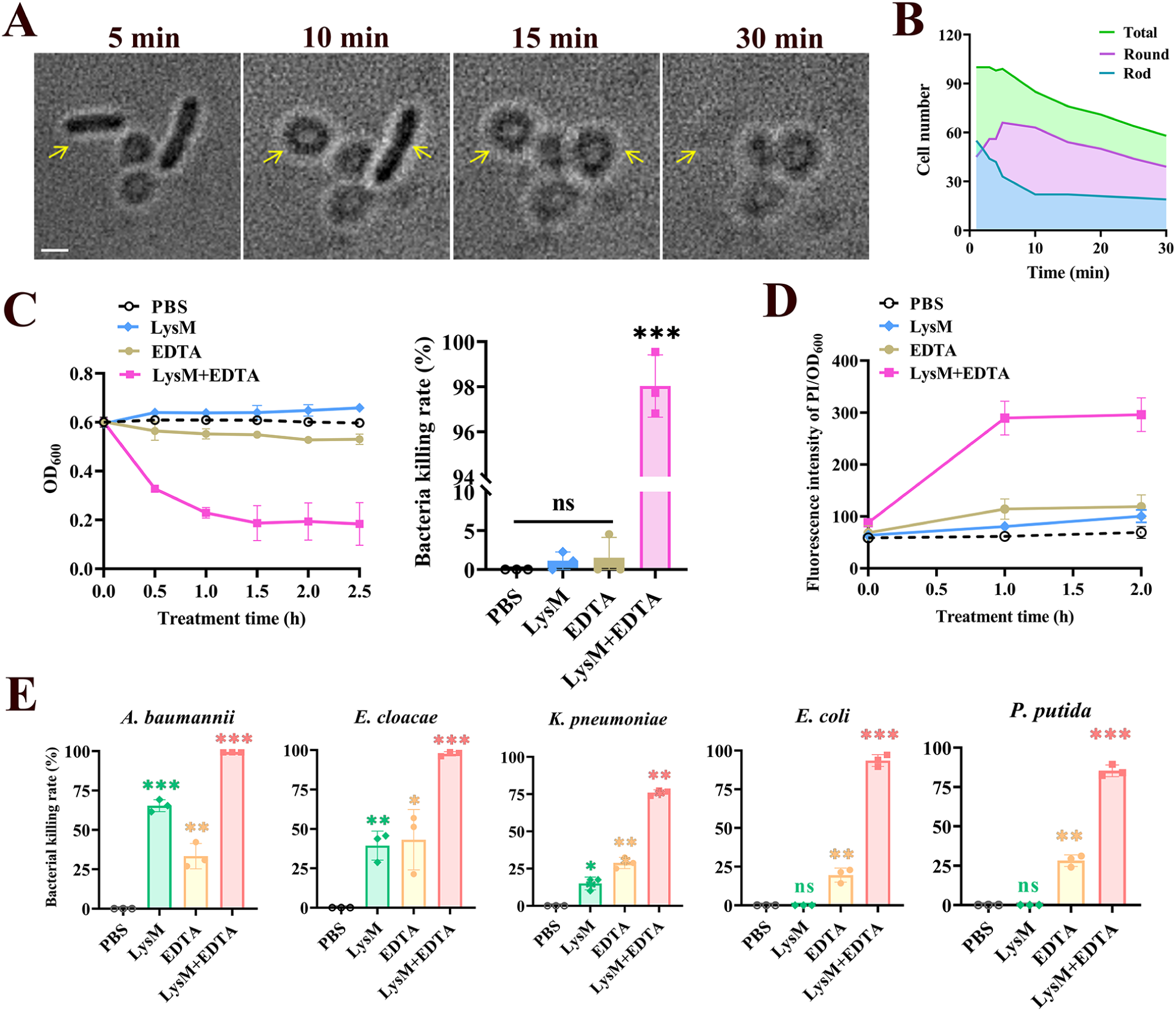
LysM exhibits bactericidal activity and leads to explosive cell lysis when supplied exogenously with EDTA. **A.** Tracking the cell morphology of *P. aeruginosa* PAO1 with the time-lapse microscopy during the treatment with LysM (30 μg/mL) and EDTA (0.5 mM) in PBS buffer. Yellow arrows indicated bacterial cells that changed the shape from rod to round. Bar, 1μm. **B.** Number of PAO1 cells with the shape of round or rod during LysM and EDTA treatment. **C.** Bactericidal effects of LysM and/or EDTA on planktonic bacteria of PAO1. Shown were the OD_600 d_uring treatments (left panel) and corresponding bacteria killing rate post 2 h treatment (right panel). **D.** Monitoring the dead bacteria during the treatment of LysM and/or EDTA on PAO1. Shown were the red fluorescence intensity of propidium iodide (PI). **E.** Bactericidal effects of LysM and/or EDTA on *A. baumannii*, *E. cloacae*, *K. pneumoniae*, *E. coli*, and *P. putida.* The corresponding bacterial killing rate (%) was shown. Error bars indicate the mean±SD of three independent experiments. Statistical significance was determined using Student’s *t-*test (*** *P* < 0.001, ** *P* < 0.01, * *P* < 0.05, ns, non-significant).

### LysM targets peptidoglycan of Gram-negative bacteria

We next examined the activity of LysM to lyse bacterial cell wall of *P. aeruginosa* in a chloroform-based turbidity assay. Results showed that the lytic activity of LysM was in a concentration-dependent manner, higher concentrations induced rapid reduction of OD_600 w_ithin the first 10 min, achieving approximately 90% of the maximal reduction by 30 min, and 50 μg/mL was found to be the most effective (Fig. 2A), which was consistent with its bacteria killing ability shown above (Fig. 1C). LysM lysed the cell wall of *A. baumannii* and *E. cloacae* with high efficiency, which was even greater than that observed for *P. aeruginosa* (Fig. 2B). In contrast, it exhibited only modest activity against *P. putida* and *E. coli* (less than 20% lysis), and showed limited lytic capacity to the cell wall of *S. aureus* (Gram-positive bacterium) regardless of chloroform pretreatment (Fig. 2B and data not shown). These results suggested that LysM was a cell wall degradation enzyme, specially targeting the peptidoglycan (PG) of Gram-negative bacteria. We then evaluated the activity of LysM to degrade commercial PG product. The PG degradation activity of LysM also exhibited a concentration-dependent manner, 30 min treatment with 50 μg/mL LysM resulted in approximately 80% PG degradation rate (Fig. 2C). The stability of LysM was evaluated across a range of pH (2–12) and temperature conditions. Maximal PG degradation activity was observed at pH 7.0 (Fig. S2A). Although relative lytic activity gradually declined as pH decreased from 7.0 to 2.0 or increased from 7.0 to 12.0, approximately 40% of the activity was retained even after exposure to pH 2.0 or 12.0. Furthermore, LysM maintained its relative enzymatic activity without significant loss after one year of storage at -80 °C or -20 °C (Fig. S2B). In summary, LysM exhibits a broad-spectrum antimicrobial activity by targeting PG, along with considerable stability across a wide range of storage and application conditions.

**Fig. 2.**
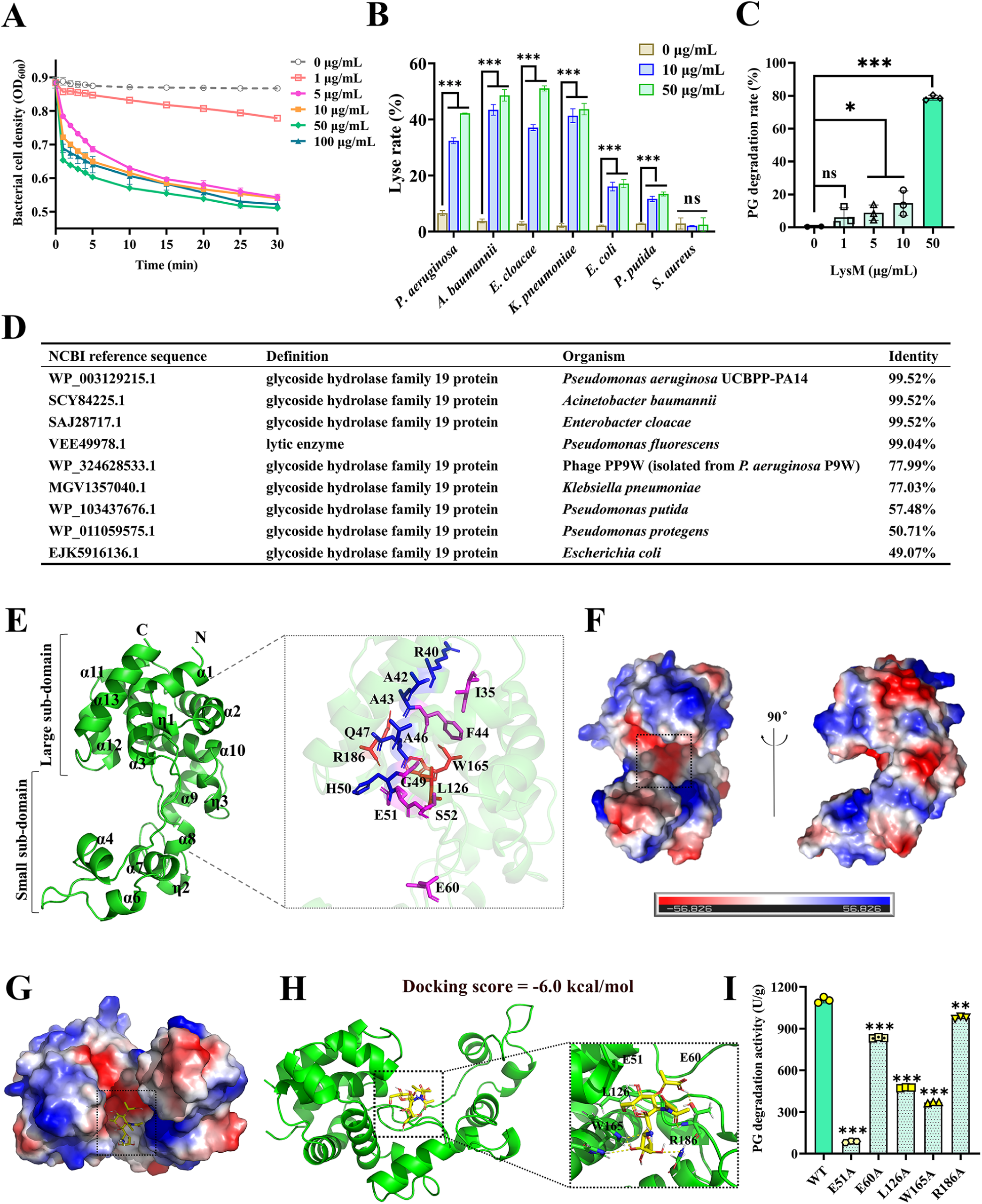
LysM targets on peptidoglycan of Gram-negative bacteria. **A.** Lytic activity of LysM against CHCl_3-_treated *P. aeruginosa* PAO1. The turbidity of PAO1 suspension was determined in the presence of LysM (0-100 μg/mL). **B.** Lytic activity of LysM against CHCl_3-_treated bacteria, namely *A. baumannii*, *E. cloacae*, *K. pneumoniae*, *E. coli*, *P. putida*, and *S. aureus*. The bacterial lysis rate post 30 min treatment was shown. **C.** The peptidoglycan degradation activity of LysM at different concentrations. **D.** Homologues of LysM with high identity were found in many bacteria and bacterial phages. **E.** The AlphaFold2-predicted 3D structure of LysM was presented in cartoon. Highly conserved residues were shown in blue or purple. W165 and R186 were shown in red. **F-G.** Electrostatic potential of LysM. The regions with negative or positive charge are colored in red or blue, respectively. The possible binding groove for GlcN was marked by dash line in G. **H.** Molecular docking diagram showed how glucosamine (GlcN) bound to LysM. The possible conformation of LysM with GlcN, and the key catalytic residues of LysM were shown. **I.** The PG degradation activity of LysM mutants, namely E51A, E60A, L126A, W165A, and R186A. Error bars indicate the mean±SD of three independent experiments. Statistical significance was determined using Student’s *t-*test (*** *P* < 0.001, ** *P* < 0.01, * *P* < 0.05, ns, non-significant).

### 3D structural modelling and molecular docking lead to the determination of key catalytic residues of LysM and its endolytic muramidase activity

InterPro sequence analysis indicated LysM (UniProt ID: G3XD75) as a single domain protein, which belongs to the CAZy <u>G</u>lycoside <u>H</u>ydrolase family 19 (GH19) (corresponding COG classification number is COG3179; E-value 1.22e-69). Enzymes of the GH19 family usually have endolytic muramidase or chitinase activity. Strikingly, LysM homologs with high sequence identity (with only one single amino acid difference) were found in *P. aeruginosa* strain PA14, *A. baumannii*, and *E. cloacae* (Fig. 2D, Fig. S3). Proteins with high similarity to LysM were distributed across many bacteria and bacterial phages, including *K. pneumoniae*, *E. coli, Pseudomonas* specie*s,* and phage PP9W, etc. (Fig. 2D, Fig. S3) and the N-terminal of LysM was highly conserved (Fig. S3B).

We then predicted the three-dimension (3D) structure of LysM using AlphaFold2 modeling, which displayed a compact “all-α-helixes” structure (Fig. 2E). The surface model of LysM (Fig. 2F) showed a negatively charged groove that was presumed to be involved in substrate binding. Highly conserved amino acid residues (I35, R40, A42, A43, F44, A46, Q47, G49, H50, E51, S52, and E60) were mostly located on α-helix 3 and near the groove region (Fig. 2E and Fig. S3B). Subsequently, we utilized this structure for molecular docking with acetylglucosamine (GlcNAc) and acetylmuramic acid (MurNAc) respectively, which yielded a favorable binding score of -6.0 kcal/mol for the LysM-GlcNAc complex. The complex positioned the GlcNAc molecule within the conserved groove (Fig. 2G-H) and suggested several key binding residues involved in substrate recognition, namely E51, E60, L126, W165, and R186 (Fig. 2E and H).

Mutations on these residues all affected the PG-degrading activity of LysM with little effect on its 3D structure based on AlphaFold2 modeling (Fig. S2C), especially E51A, L126A, and W165A mutations reduced the PG-degrading activity by 97%, 66% and 73%, respectively, indicating their critical role (Fig. 2I). Given the PG degrading activity of LysM and the fact of LysM: i) cannot degrade chitin (data not shown); ii) strongly interacts with GlcNAc, but not with MurNAc in docking analysis; iii) shows high similarity to the recently reported PaGH19Lys with muramidase activity(30); we proposed that LysM was an endolytic muramidase.

### LysM in medium can inhibit biofilm formation of *P. aeruginosa* by interfering with bacterial growth and surface adherence

As shown above, exogenous LysM requires outer membrane disruption in order to enter periplasmic space to hydrolyze PG. We next investigated whether LysM alone (without EDTA to disrupt outer membrane) has anti-biofilm activity against *P. aeruginosa* in growth media. Strikingly, LysM in media was able to completely inhibit biofilm formation of *P. aeruginosa* and its minimum biofilm inhibitory concentration (MBIC) was 6.25 μg/mL (Fig. 3A). The biofilm inhibition of LysM was likely due to an inhibition on bacterial growth because the minimum inhibitory concentration (MIC) of LysM in medium was also 6.25 μg/mL (Fig. 3B). The addition of EDTA in media did not enhance the inhibition activity of LysM on biofilm or bacterial growth (Fig. 3A-B and Fig. S4B-C).

**Fig. 3.**
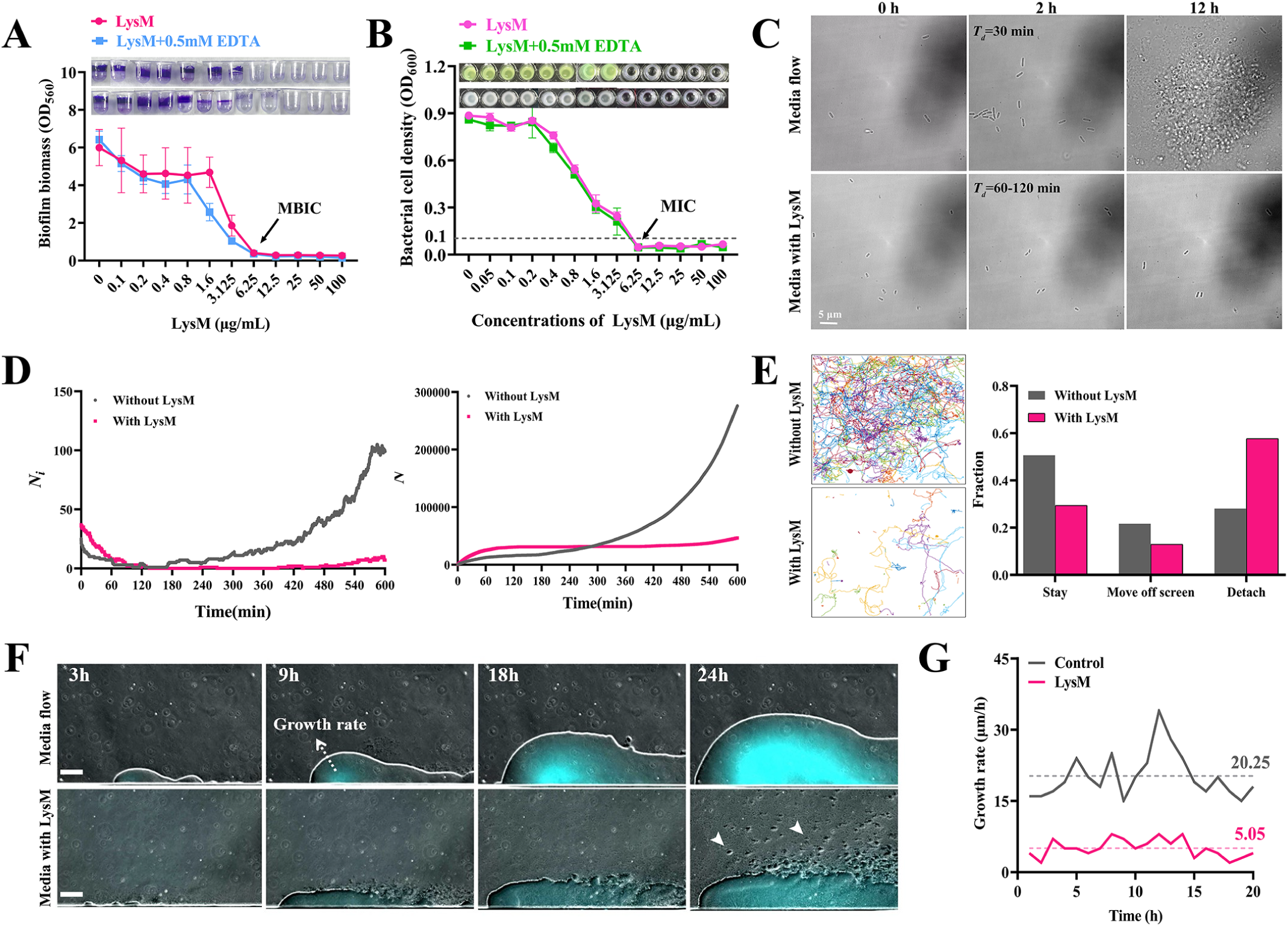
LysM alone can inhibit the growth and biofilm formation of *P. aeruginosa* in growth medium. **A.** The inhibitory effects of LysM with or without EDTA on biofilm formation of *P. aeruginosa* PAO1. Arrow indicated the corresponding minimal biofilm inhibitory concentration (MBIC). The biofilm biomass was measured by CV assay. **B.** The inhibitory effects of LysM with or without EDTA on the growth of PAO1 and the arrow indicated the minimal inhibitory concentration (MIC)**. C-E.** The impact of LysM on PAO1 biofilm formed in the flow-cell system. Shown were bacteria attached on the surface of the flow-cell at time that the treatment was initiated (0 h) and after the cultivation of 2 h and 12 h (C), the number of surface-adhered bacteria at the time point of image acquired (D, Left panel), the cumulative number of surface-adhered bacteria during 12h tracking (D, Right panel), trajectory of surface-attached bacteria within 12 h tracking (E, left panel, each color indicates the trail of a tracked bacterial cell), the percentage of bacteria that are stable attached on surface (stay), move off the screen, and detached from the surface during the tracking (E, right panel). **F-G.** Effects of LysM on the *P. aeruginosa* biofilm grown in a microfluidic device with continuous medium supply. Shown were time-lapse images of biofilms of PAO1 during 24 h growth, which were a merge of phase contrast and fluorescent images. The fluorescence (shown in cyan) was the signal of pyoverdine produced by PAO1. The short arrows indicated the bacterial aggregates. The corresponding growth rate of biofilm was shown in G.

In order to know how LysM inhibits biofilm formation, we tracked biofilm formation in a flow-cell system with or without addition of LysM in medium. Several surface-attached bacteria were tracked for 12 h (Fig. 3C, Video 3). Without LysM, PAO1 can formed a 3D biofilm post 12 h growth in a flow-cell. Remarkably, the addition of LysM completely blocked biofilm formation of PAO1 (Fig. 3C). During the first 2 h-LysM treatment, bacterial cells were live and able to divide, but with a relatively slower growth rate (the doubling time was approximately twice of untreated bacterial cells) (Fig. 3C, Video 2A-B); more importantly, one of the daughter bacterial cells tended to detach from the surface and flow away (Video 2A and 2B). The phenomenon of slow growth was similar to what has been reported for the strain with mutation on cell wall-hydrolase (31). This suggested that LysM might affect the bacterial growth by interfering the remodeling of peptidoglycan during cell division, since LysM is a <u>g</u>lycoside hydrolase that hydrolyses PG. Analysis of all images acquired during 12-h of tracking indicated that the number of surface-attached bacteria in the flow-cell with LysM treatment remained very low throughout the 12-h period, whereas in the media control flow-cell, the number of surface-attached bacteria increased exponentially (Fig. 3D). In addition, LysM increased the percentage of bacteria that detached from the surface (Fig. 3E, right panel, detach) and decreased the number of bacteria stably attached on the surface (Fig. 3E, right panel, stay). This was consistent with much fewer moving trails of surface-adhered bacteria in the LysM treatment group (Fig. 3E, left panel). Taken together, our data showed LysM inhibited the biofilm formation of *P. aeruginosa* in growth media by interfering bacterial growth and stable surface adherence.

We then further investigate the inhibition of LysM on biofilm formation of PAO1 in a microfluidic device designed by Liu et al.(32), in which the regular 3D biofilm was restricted in a semi-2D biofilm structure and initiated by a small microcolony at inoculation. During the 24 h tracking (Fig. 3F and 3G, Video 3A and 3B), the biofilm expansion of PAO1 was seriously limited by LysM, resulting in a significantly lower growth rate (5.05 μm/h), which is 4-fold less than that of untreated control (20.25 μm/h) (Fig. 3G). The expanding edge of untreated biofilm was sharp and smooth (Fig. 3F, Video 3A), whereas the LysM-treated biofilm had a relatively irregular and rough edge, with visible bacterial cells and aggregates in the media flow (Fig. 3F, indicated by the short arrows) suggesting that biofilm was dispersing during treatment (Fig. 3F, Video 3B). This result further indicated that LysM alone was able to restrict the growth of biofilm in medium.

### The inhibition of LysM on other Gram-negative bacteria and their biofilm in media

Addition of LysM in media did show significant inhibition on the growth and biofilm formation of *A. baumannii* and *E. cloacae* with less efficacy than that of *P. aeruginosa*, but LysM alone was not able to inhibit the growth and biofilm formation of *K. pneumoniae, E. coli*, or *P. putida* (Fig. 4A and 4B). However, the addition of EDTA with LysM did promote their inhibition on all tested strains (Fig. 4A and 4B). We then further examine the effect of LysM with EDTA on a preformed *E. coli* biofilm in microtiter dish and the microfluidic device. When a 20 h-old *E. coli* biofilm was exposed to LB supplemented with LysM or in combination with EDTA for 2 hours, LysM with EDTA reduced the half of biofilm biomass and killed 2 log of biofilm bacteria (Fig. S4D). For the 20 h-old biofilm grown in the microfluidic device, LysM with EDTA was added to the LB medium and treated for another 20-h. The untreated biofilm gradually expanded with an average growth rate of 27.8 μm/h. While the addition of LysM and EDTA dramatically limited biofilm expansion with a continuously decreasing growth rate, the average growth rate was only 4.8 μm/h (Fig. 4C and 4D, Video 4A and 4B). To track the cell death during the growth of biofilm, PI was also added to the media at inoculation. Post 20 h growth, no detectable red fluorescent signal was found, suggesting little cell death occurring within the biofilms (Fig. 4C, left panel). After LysM and EDTA applied, the red fluorescent signal rapidly increased approximately 150-fold during 20 h treatment, while the red fluorescence of the untreated biofilm was 12-fold lower (Fig. 4C and 4E). These results demonstrated that LysM and EDTA effectively suppressed the growth of a formed-biofilm and killed bacterial cells within biofilm under continuous medium supply condition.

**Fig. 4.**
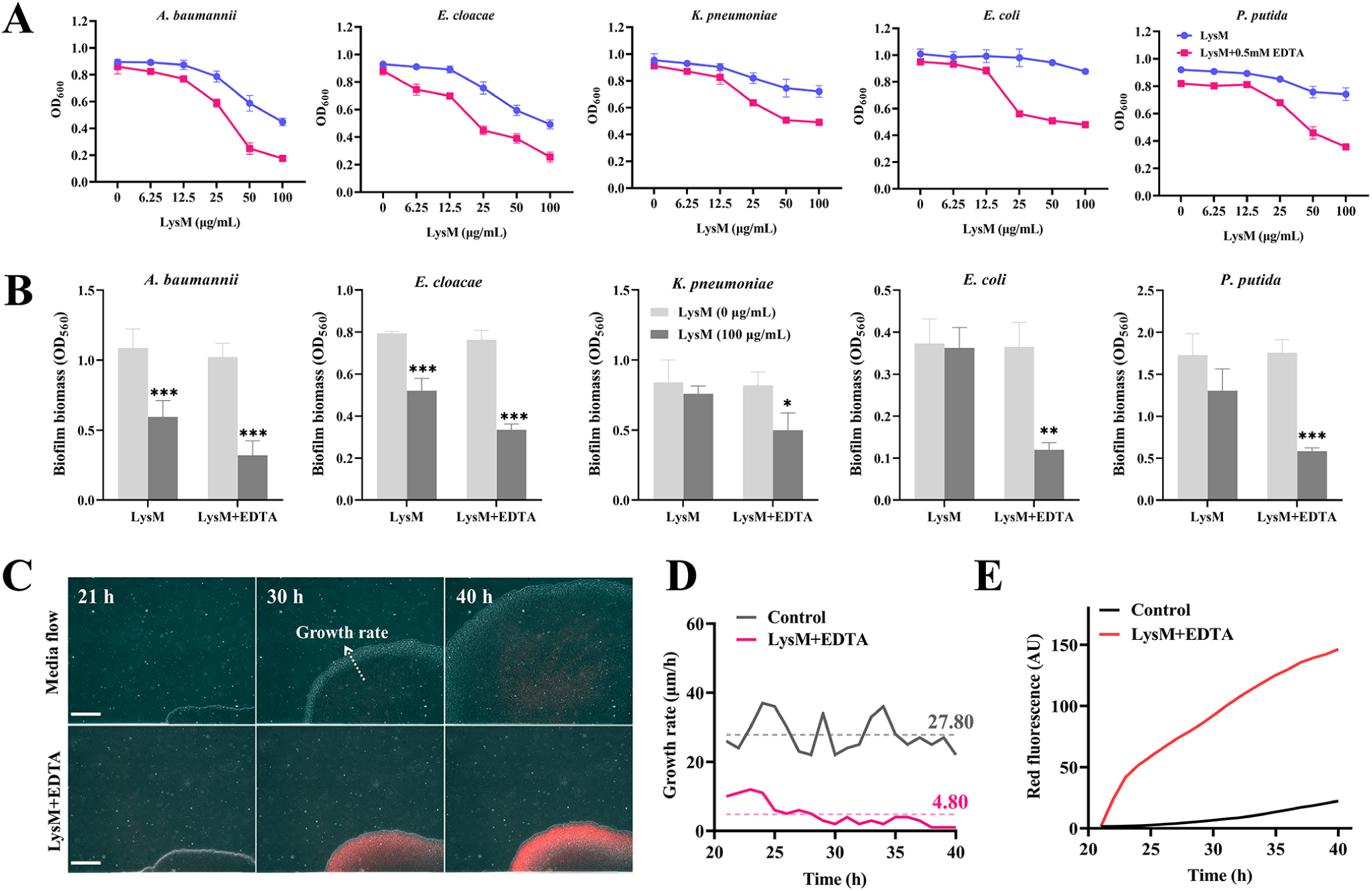
The inhibitory effects of LysM alone or with EDTA on the growth and biofilm formation of *A. baumannii*, *E. cloacae*, *K. pneumoniae*, *E. coli*, and *P. putida* in media. A. The inhibitory effects of LysM alone or with EDTA on bacterial growth in corresponding growth media. B. The inhibitory effects of LysM alone or with EDTA on biofilm formation of indicated bacteria. C-E. Inhibitory effect of LysM (30 μg/ml) and EDTA on *E. coli* biofilm grown in the microfluidic device under continuous medium supply. The time-lapse images (merge of phase contrast and red fluorescence) were shown in (C), the biofilm growth rate was shown in (D), and the changes of total red fluorescent intensity during the 40h-growth were shown in (E). The red fluorescent signal indicated the PI-stained dead bacteria within biofilms. Scale bar, 100 μm. Error bars indicate the mean±SD of three independent experiments. Statistical significance was determined using Student’s *t-*test (*** *P* < 0.001, ** *P* < 0.01, * *P* < 0.05, compared to the untreated sample).

### The LysM-based “cocktail” efficiently disrupts and eliminates 24 h-old biofilms formed by nosocomial pathogens and MDR clinical isolates of *P. aeruginosa*

Many antibiotics cannot efficiently kill biofilm bacteria. Therefore, we investigated the killing effect of LysM with EDTA on biofilms. LysM ranging from 6.25 to 50 μg/mL was applied on biofilms of PAO1. Consistent with the above bacterial killing rate, LysM (over 6.25 μg/mL) combined with EDTA significantly decreased the biofilm biomass of a 24 h biofilm within 2 h, while the EDTA or LysM (6.25 to 50 μg/mL) alone failed to destroy the established biofilm in PBS buffer (Fig. 5A). Viable bacterial counts indicated that LysM with EDTA efficiently killed biofilm bacteria and the killing rate was higher than 90% (Fig. 5B). Consistent with the killing rate of planktonic bacteria (Fig.1C), LysM with EDTA exhibited biofilm bacterial killing rate of approximately 90-100% after 2 h of treatment against the other Gram-negative ESKAPE bacteria, *A. baumannii*, *E. cloacae*, and *K. pneumoniae* as well as *E. coli* (Fig. 5F and Fig. S5). These results indicated that LysM with EDTA was able to eliminate biofilms formed by Gram-negative pathogens in the ESKAPE group.

**Fig. 5.**
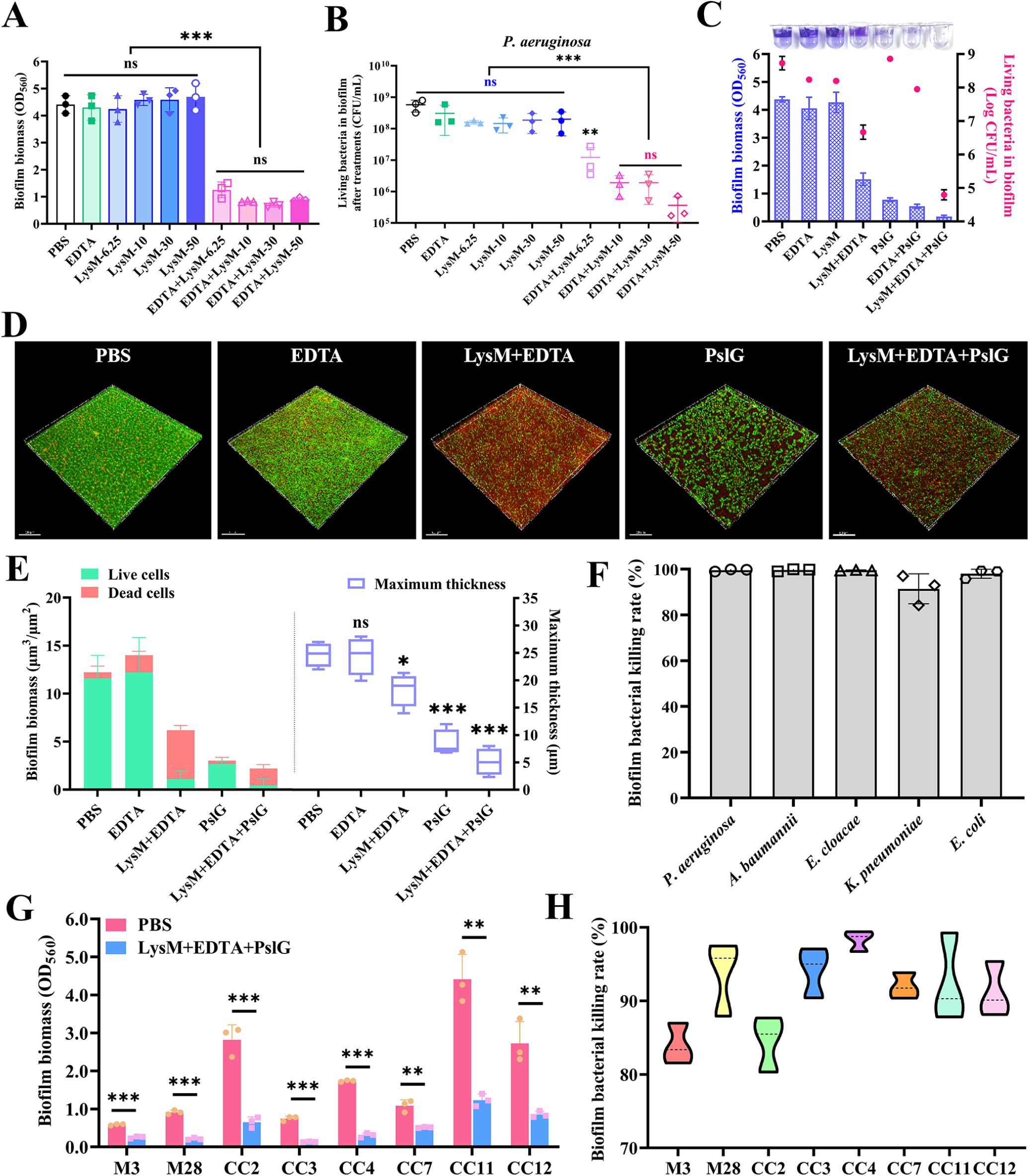
The LysM-based “cocktail” efficiently disrupts and eliminates biofilms of multiple Gram-negative pathogens. **A-B**. The effect of LysM on 24 h-old biofilm of *P. aeruginosa* PAO1. Biofilm biomass (A) and living bacteria within biofilm (B) were examined after treatments of LysM with or without EDTA in PBS buffer at 37 °C for 2 h in microtiter dish. **C.** Effects of LysM “cocktail” on the elimination of PAO1 biofilm in microtiter dish. Show was biofilm biomass and living bacteria within biofilm post 2 h treatment. The wells post crystal violet staining were shown above corresponding column. **D-E.** The impacts of LysM-based “cocktail” on pellicles of PAO1. Confocal laser scanning microscopy (CLSM) images of pellicles post treatment were shown in D. PAO1 pellicles were treated with LysM and/or ETDA and/or PslG for 2 h, following the live(green)/dead(red) staining by SYTO9 and PI. Shown in E were the corresponding biofilm biomass and the maximum thickness quantified from CLSM images. **F.** The biofilm bacterial killing rate (%) of the LysM+EDTA treatment for 2 h against *A. baumannii*, *E. cloacae*, *K. pneumoniae*, *E. coli*, and *P. aeruginosa.* **G-H**. Anti-biofilm activity of LysM “cocktail” against antibiotic resistant clinical *P. aeruginosa* isolates, namely M3, M28, CC2, CC3, CC4, CC7, CC11, and CC12. The biofilm biomass before and after treatment was shown in (G), and the biofilm bacterial killing rate (%) was shown in (H). Error bars indicate the mean ± SD of three independent experiments. Error bars indicate the mean±SD of three independent experiments. Statistical significance was determined using Student’s *t-*test (*** *P* < 0.001, ns, non-significant).

Our previous work has shown that PslG (a glycosyl hydrolase that degrades biofilm matrix exopolysaccharide Psl) sensitizes biofilm bacteria to antimicrobial agent (27). Therefore, we further investigated the efficacy of LysM+EDTA+PslG “cocktail” on biofilm eradication. PslG indeed further enhanced the killing effect of LysM+EDTA on biofilms formed by *P. aeruginosa* PAO1 strain. The “cocktail” greatly improved the eradication of biofilm, destroying more than 95% of biofilm biomass in a microtiter dish biofilm assay (Fig. 5C), and it also significantly decreased the viable cells by more than 4 log (Fig. 5C), indicating that 99.99% of biofilm bacteria were killed. This efficacy was comparable to the combinations of PslG with 2× MIC of CIP or TOB (Fig. S4A). The LysM-based “cocktail” was further examined on its effect on pellicles, the air-liquid interface biofilms. Live/dead staining results showed that LysM+EDTA treatment reduced the half of pellicle biomass during 2 h and killed 90% live bacterial cells compared to PBS treatment control (Fig. 5D-E), while the LysM+EDTA+PslG “cocktail” eliminated 99% of biofilm bacteria and disrupted the 82% of pellicle biomass (Fig. 5D-E).

We then further examined whether this “cocktail” has the potential as an alternative therapy to effectively fight against biofilms formed by MDR clinical *P. aeruginosa* isolates. Eight MDR clinical isolates (Table S2) were selected for the biofilm elimination assay. The LysM “cocktail” showed biofilm clearance rate ranging from 50% to 80% for the MDR isolates and corresponding biofilm bacterial killing rate was all higher than 80% (Fig. 5G-H). Taken together, our results suggested that the LysM-based “cocktail” had great potential for biofilm eradication to combat nosocomial infections.

### Evaluation of LysM and its “cocktail” for combating biofilm-related *P. aeruginosa* infection using human airway organoids (HAO) model

Given that LysM and its cocktail had minimal effect on the viability of the human lung epithelial A549 cells (Fig. S6) and their promising biofilm elimination activity, especially on biofilms formed by *P. aeruginosa*, we thus further evaluated the biofilm inhibition and eradication activity of LysM using *P. aeruginosa* human airway organoids (HAO) infection model. In the biofilm inhibition experiments, eight different treatments were applied at the time of PAO1 inoculation. At 24 h post-infection, biofilms were detected and quantified by <u>c</u>onfocal laser <u>s</u>canning <u>m</u>icroscopy (CLSM) and the HAOs were examined by hematoxylin and eosin (H&E) staining, immunofluorescence, and transepithelial electrical resistance (TEER). Consistent with the *in vitro* data, LysM alone significantly inhibited the biofilm formation on HAOs. However, the LysM+EDTA+PslG “cocktail” showed the best biofilm inhibition efficacy, which was comparable to that of the PslG+CIP treatment (Fig. 6A and 6B). Beyond bacterial reduction, the “cocktail” treatment demonstrated superior protection of the organoid structure. The results of H&E staining and immunofluorescence both revealed that the “cocktail” maintained a highly organized pseudostratified epithelial architecture, while the PBS and EDTA groups suffered from severe epithelial sloughing and loss of structural continuity, which was indicated by the columnar morphology of epithelial cells, the integrity of the apical cilia, and the distribution of goblet cells in HAOs. Furthermore, TEER measurements also indicated that the LysM+EDTA+PslG “cocktail” preserved the epithelial barrier function to the greatest extent (Fig. 6A and 6B, Fig. S7).

**Fig. 6.**
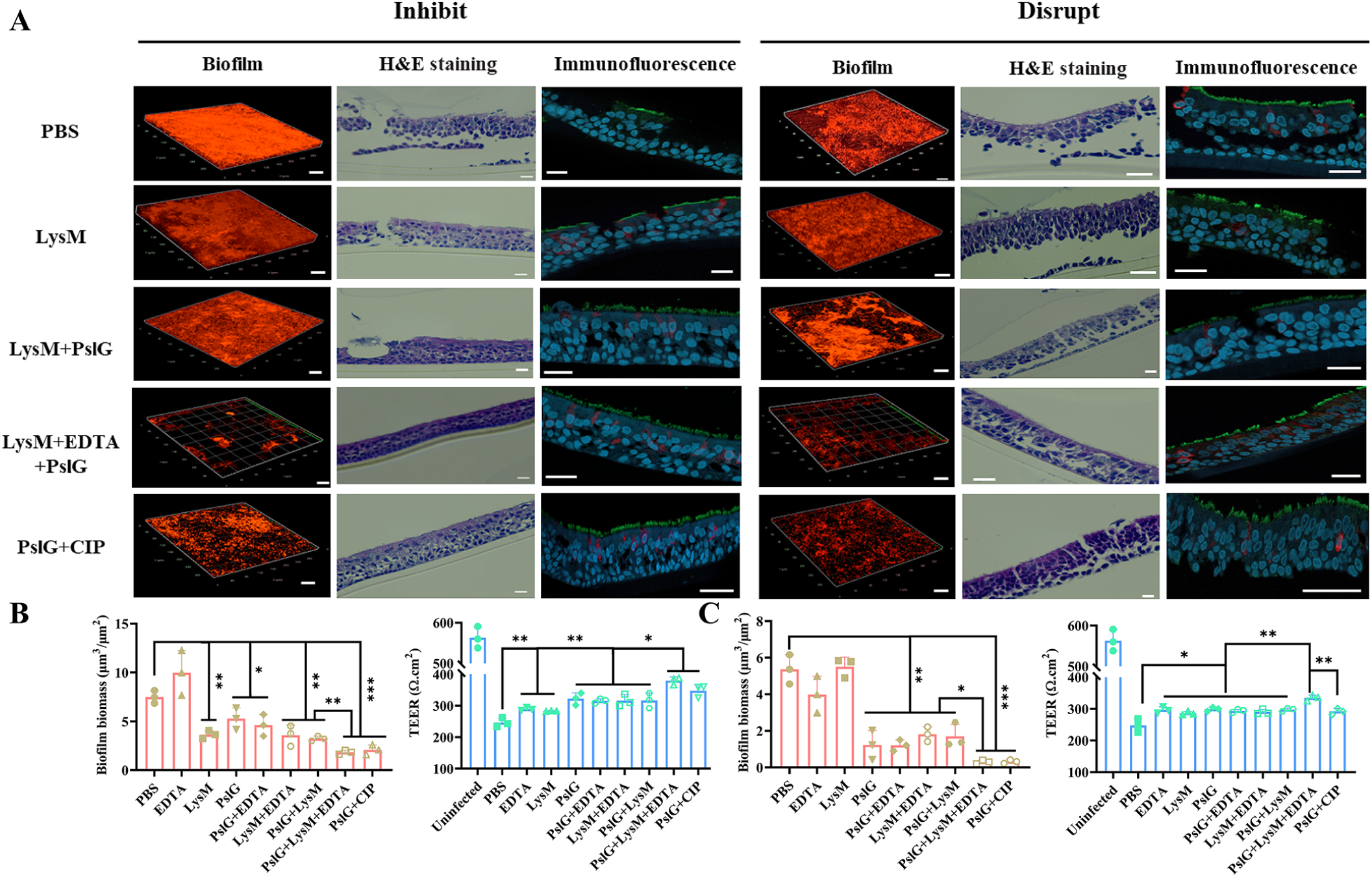
LysM “cocktail” inhibits biofilm formation and disrupts the 24 h-old biofilm of *P. aeruginosa* grown on human airway organoids (HAOs). A. Visualization of biofilm architecture, tissue/cellular morphology, and spatial distribution of mucin of human airway organoids (HAOs) under different treatment groups, namely PBS buffer, 30 μg/mL LysM in PBS buffer, LysM+PslG (PBS buffer containing 10 μg/mL LysM and 50 nM PslG), LysM+EDTA+PslG (PBS buffer containing 10 μg/mL LysM, 0.5 mM EDTA, and 50 nM PslG), and PslG+CIP (PBS buffer containing 50 nM PslG and 1 μg/mL CIP). *P. aeruginosa* PAO1::*mScarlet-I* strain was applied. For biofilm inhibition assay, 10^5^ bacteria in PBS buffer with different treatments were inoculated onto HAO. After 24 h of growth, the biofilms were washed three times with PBS and imaged by CLSM. Thereafter, the corresponding HAOs were fixed and sectioned. The HAOs sections were assessed by H&E staining and immunofluorescence. For biofilm disruption assay, the 24 h-old biofilms grown on HAOs were washed with PBS, applied the various treatments for 6h, and then imaged by CLSM. For immunofluorescent staining, the cilia of HAO stained in green were detected by anti-acetyl-α-tubulin antibody, mucin stained in red was marketed by anti-MUC5AC antibody, and the nuclei of epithelial cells were counterstained by DAPI (blue). A reprehensive image of each treatment group was shown. Scale bars = 10 μm. **B-C.** Biofilm biomass quantified from CLSM images and TEER (transepithelial electrical resistance) of corresponding HAOs indicating the effects of treatments on biofilm and epithelial barrier integrity. The results of biofilm inhibition (B) and disruption (C) were shown. Error bars indicate the mean±SD of three independent experiments. Statistical significance was determined using Student’s *t-*test (*** *P* < 0.001, ** *P* < 0.01, * *P* < 0.05, ns, non-significant).

In the biofilm eradication experiment, PAO1 biofilms were established on HAOs for 24 h post-inoculation, followed by various treatments for 6 h. Thereafter, biofilms and HAOs were examined. Notably, LysM+EDTA+PslG and PslG+CIP treatment showed the best biofilm eradication, the other treatments, except for EDTA and LysM, all exhibited significant biofilm disruption (Fig. 6A and 6C, Fig. S7). Additionally, H&E staining, immunofluorescence, and TEER results indicated that the LysM+EDTA+PslG combination was more effective than the PslG+CIP combination in regard to preserving the integrity of tight junctions between HAO cells (Fig. 6A and 6C, Fig. S7).

In summary, our results indicated that the LysM+EDTA+PslG “cocktail” had significant efficacy in inhibiting and eradicating *P. aeruginosa* biofilm in a HAO infection model, highlighting its potential as an alternative to traditional antibiotics.

### LysM enhances the efficacy of antibiotics against multiple Gram-negative nosocomial pathogens

To know whether LysM might have synergistic activity in combination with antibiotics against *P. aeruginosa* related infections. We performed a chequerboard assay to determine the potency of LysM in combination with several antibiotics that are commonly used in clinic. Notably, no antagonism was observed across all the antibiotics tested in the study. Using PAO1 as a testing strain, the synergy was observed between LysM with CIP (FICi = 0.281), TOB (FICi = 0.375), ERY (FICi = 0.375) and CRB (FICi = 0.375), while additive effects (0.5 < FICi ≤ 1.0) were found for LysM combined with TET or colistin (Fig. 7A). We further examined whether LysM can reverse the sensitivity of MDR clinical isolates of *P. aeruginosa*. Two MDR strains, CC7 and CC11 were selected for the assay, both of which are carbapenem-resistant isolates (Table S2). LysM reduced the MIC of both CIP and colistin for 2-fold against the two MDR strains respectively (Fig. S8A). To evaluate the broad potentiating spectrum of LysM, we determined the effect of LysM on MICs of CIP and colistin against other ESKAPE pathogens, *A. baumannii*, *E. cloacae*, and *K. pneumoniae*, respectively. The addition of LysM significantly reduced the MICs of both antibiotics against all three species (Fig. 7B-C). These data showed that LysM could be also used as an antibiotic adjuvant to enhance antibiotic sensitivity of pathogens in the EKAPE group.

**Fig. 7.**
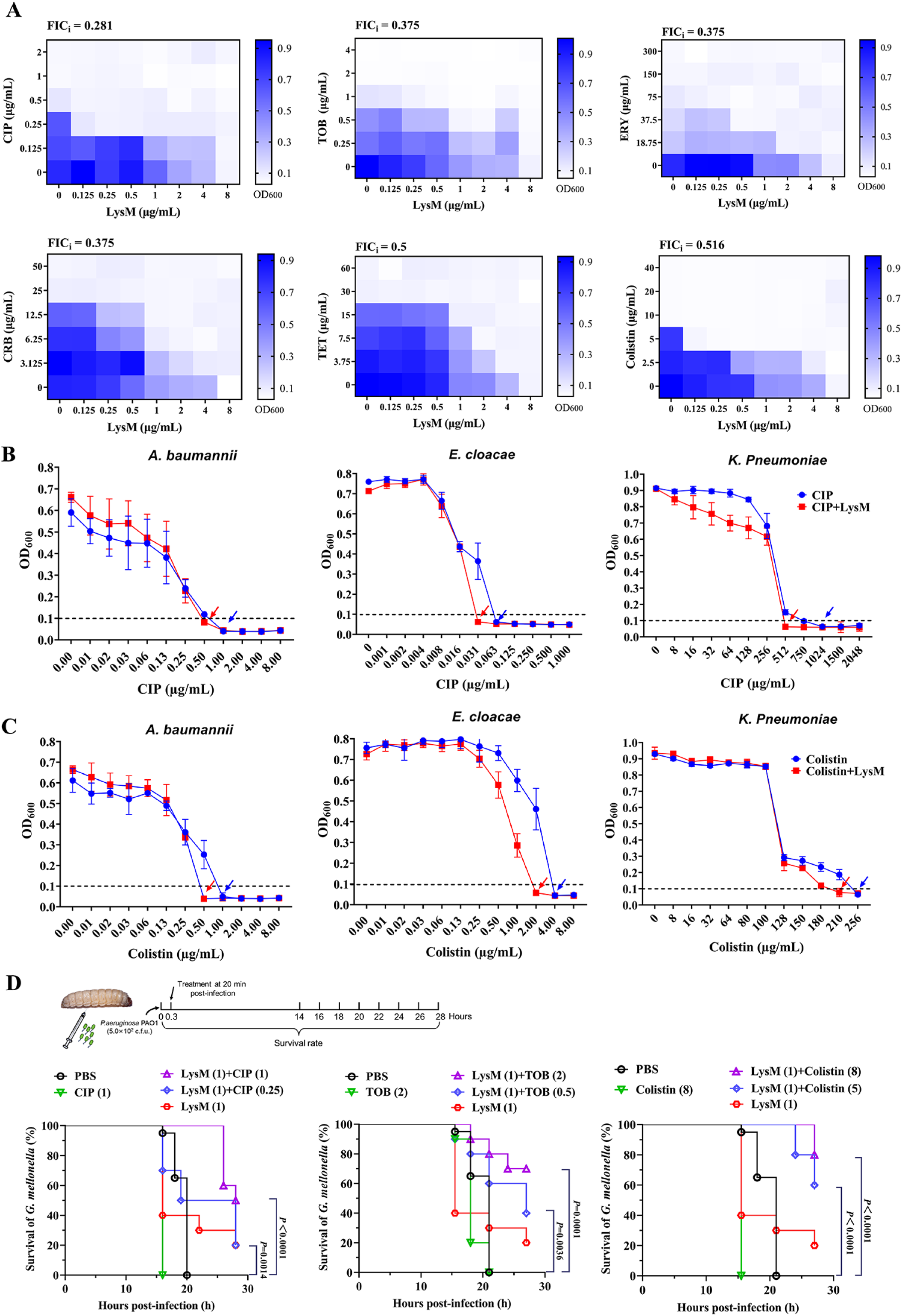
LysM sensitizes Gram-negative ESKAPE pathogens to antibiotics. **A.** Chequerboard broth microdilution assays of LysM with CIP, TOB, ERY, CRB, TET and Colistin against *P. aeruginosa* PAO1 respectively. The interactions were interpreted based on the calculation of FICi, in which: FICi ≤ 0.5, synergistic; 0.5 < FICi ≤ 1, additive; 1 < FICi ≤ 2, neutral; and FICi ⩽ 2, antagonistic, respectively. **B-C.** The impact of LysM (10 μg/mL) on the MIC value of CIP (B) and Colistin (C) against *A. baumannii*, *E. cloacae,* and *K. pneumoniae*. The solid arrows indicate the MIC. D. *In vivo* therapeutic efficacy of LysM-antibiotic combinations in a *G. mellonella* infection model. Top: Schematic of the experimental design. Bottom: Kaplan-Meier survival curves of *G. mellonella* larvae infected with a lethal dose of PAO1 (5.0 × 10^2^ c.f.u.) and treated 20 min post-infection with LysM (1 μg/mL), antibiotics (1 μg/mL CIP, 2 μg/mL TOB, or 8 μg/mL Colistin), or their respective combinations. Statistical significance is indicated by the respective *P* values.

We next evaluated the *in vivo* therapeutic efficacy of LysM-antibiotic combinations using a *Galleria mellonella* killing model infected with *P. aeruginosa* PAO1 (Fig. 7D, Fig. S8B). At 28 h post infection, the larvae treated with the combination of LysM with either CIP, TOB, or colistin, all showed strikingly increased survival rate. Overall, our findings collectively demonstrated the potential of LysM as a promising antimicrobial agent or antibiotic adjuvant against nosocomial infections.

## Discussion

The rapid emergence of antibiotic resistance in bacteria is a global challenge. The tolerance and resistance of microbial biofilms to antimicrobials and to the immune system are a main cause of persistence of chronic infections and also one of the major sources of MDR bacteria. Therefore, new therapeutic strategies and antimicrobials are urgently required to combat pathogens, especially biofilm-forming and antibiotic-resistant pathogens. In this study, we have shown LysM, a glycoside hydrolase that specially targets PG of Gram-negative bacteria, can efficiently kill planktonic and biofilm of Gram-negative bacteria in ESKAPE group when applied with EDTA as a bacterial outer membrane permeabilizer.

LysM was encoded by PA0629 located on the *P. aeruginosa* genome. Its homologues can be found in many bacterial genomes as well as phages. Interestingly, LysM has shown the best antimicrobial and anti-biofilm activity against *P. aeruginosa* among the ESKAPE group pathogens. More remarkably, LysM alone inhibits the growth and biofilm formation of *P. aeruginosa* at a low concentration, but shows much lower inhibition effect on other bacterial species. The 3D structure of LysM and its key catalytic residues that we have reported in this study may help to define the associated mechanisms of these specific actions in the future.

*P. aeruginosa* infections are difficult to eradicate due to its ability to form biofilms and its intrinsic antibiotic resistance. In this study, we present the LysM-based “cocktails” with high efficacy of bactericidal and biofilm eradication against laboratory *P. aeruginosa* strain as well as clinical MDR isolates. In addition, LysM can function as an adjuvant to sensitize the MDR *P. aeruginosa* to common antibiotics, suggesting alternative strategies to combat clinic MDR isolates. The HAO and *G. mellonella*infection models demonstrate the potency of LysM as a clinical candidate for treatment of *P. aeruginosa* infections.

Overall, this study opens new avenues for the treatment of nosocomial infections. Our findings have shown that LysM has potent bactericidal and biofilm eradication activity along with the function as an antibiotic adjuvant, which suggest the great potential of LysM as a clinical candidate to fight against nosocomial infections. The findings also provide a scientific foundation for the development of novel antimicrobials since LysM homologues are widely distributed across bacterial and phage genomes.

## Materials and methods

### Bacteria strains and growth conditions

Bacterial strains and plasmids used in this study are listed in Table S1. The multidrug-resistant strains used in this study were isolated from the sputum samples of clinical patients, and their antibiotic resistance profile has been characterized. Unless otherwise indicated, all strains were routinely cultured in Luria-Bertani (LB) broth (Becton Dickinson) at 37 °C. For biofilm formation, *Pseudomonas aeruginosa* was cultured in Jensen’s medium, a chemically defined medium, at 30 °C, while *Acinetobacter baumannii*, *Enterobacter cloacae*, *Klebsiella pneumoniae*, *Escherichia coli*, and *Pseudomonas putida* were cultured in LB broth at 30°C. Kanamycin at 50 μg/mL was added into the medium for *E. coli* with pET28a-derived plasmids.

### Plasmid construction and protein purification

To construct the plasmid for LysM protein purification, the entire PA0629 (*lysM*) gene was amplified from the genomic DNA of *P. aeruginosa* PAO1 and cloned into pET28a (Novagen, USA) at EcoRI and HindIII sites. For the expression of LysM in *E. coli*, the recombinant plasmid pET28a-*lysM* was transformed in *E. coli* BL21(DE3), and the protein was expressed as follows: transformants were grown overnight in LB containing kanamycin at 50 μg/mL. The overnight culture was diluted and incubated to attain an OD_600 o_f 0.6. The transcription of LysM was induced by the addition of isopropyl-β-D-thiogalactopyranoside (IPTG) at 1 mM and incubation at 18 °C for 20 h (27, 33). The cells were harvested by centrifugation and suspended in 10 mL lysis buffer containing 1 mmol/L phenylmethylsulfonyl fluoride (PMSF, Nacalai Tesque, Inc., Kyoto, Japan), 20 mmol/L sodium phosphate, and 200 mmol/L NaCl (pH 8.0). The cell suspension was cooled on ice and disrupted by sonication for 8–10 cycles of 30 s pulse and 30 s pause using an ultrasonic disruptor UD201 (Tomy Seiko Co., LTD, Japan). The supernatant containing LysM was collected by centrifugation (12,000×g, 4 °C, 30 min) and purified using a HisTrap FF crude column (GE Healthcare Bio-Sciences AB, Uppsala, Sweden). The purity of the protein was confirmed by sodium dodecyl sulfate-polyacrylamide gel electrophoresis (SDS-PAGE). The concentration of LysM was determined using BCA kit.

### Time-lapse microscopy

Time-lapse microscopy was employed to continuously monitor the cell lysis in the presence of LysM and EDTA. Briefly, PAO1 grown in exponential phase was harvested and re-suspended in PBS at 10^7^ CFU/mL, LysM was added to reach 30 μg/mL, and 10 μL of this mixture was placed on the agar, which was fixed on a glass coverslip in advance. The sample was incubated at 37 °C and observed using an inverted Olympus IX83 microscope (Olympus, Japan) equipped with a 100×objective lens to capture Phase-contrast and fluorescence images were acquired every 30 s using an Andor Zyla 4.2 sCMOS camera (Andor Technology, UK), with the first image designated as 1 min post-treatment. Data analysis was conducted using ImageJ (National Institutes of Health, USA) and MATLAB (MathWorks, USA).

### Bioinformatic analysis and molecular docking

#### According to the gene prediction, one possible sequence PA0629 encoding was identified from *Pseudomonas aeruginosa* PAO1 sequences (<u>The *Pseudomonas*</u>

Genome Database - Genome annotation and comparative genome analysis), and named as *lysM.* The conserved domain architecture of LysM (encoded by PA0629) was analyzed by NCBI Conserved Domain Search (https://www.ncbi.nlm.nih.gov/Structure/cdd/wrpsb.cgi). Structure of LysM was computed with the Alphafold2 server (https://colab.research.google.com/github/sokrypton/ColabFold/blob/main/AlphaFold2.ipynb), and the highest pLDDT score was chosen as its protein model. In addition, ChimeraX software was used to demonstrate the basic structure of the protein, its surface charge, and the putative B-factor parameters. The generated model structure was visualised and analysed using the PyMOL molecular graphics system (http://www.pymol.sourceforge.net/). The docking analysis for LysM and ligand structure of glucosamine (GlcN), a monomer unit of PG, was performed using the Swissdock server (www.swissdock.ch) with default parameters according to the reported protocol. The molecular docking scores and interaction images were predicted.

### PG degrading assay

The peptidoglycan (PG) degradation ability of LysM was determined using a commercial PG product (ACMEC biochemical, China) according to the manufacturing instructions. Briefly, the PG powder was suspended in sterile water at 1 mg/mL, then 20 μL of LysM solution (0-0.5 mg/mL) was mixed with 180 μL of PG suspension in a 96-well plate. The mixture was incubated at 37 °C for 2 h, and the OD_450 w_as measured. The PG degradation rate was calculated by using the following equation:

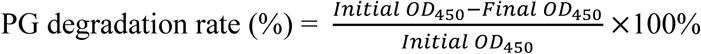

### Lysis assay of LysM

The preliminary lytic activity assay of LysM against bacterial cell wall without outer membrane was measured using the turbidity method described by with slight changes (33). Briefly, PAO1 harvested from the overnight culture were suspended in 50 mM Tris-HCl buffer (pH 7.7) saturated with chloroform (CHCl_3)_, and incubated at 37 °C for 15 min to remove bacterial outer membrane (34, 35). The pretreated cells were centrifuged, washed twice and re-suspended in PBS at OD_600 =_ 0.6–0.8. Then 20 μL of LysM solution (0-1 mg/mL) was mixed with 180 μL of CHCl_3-_treated cell suspension in a 96-well plate. The mixture was incubated at 25 °C for 30 min, and the OD_600 w_as recorded per minute. Lysis rate (%) was calculated using the following equation:

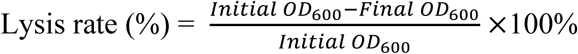

### Stability test of LysM

The stability of LysM was evaluated under various storage temperature and pH according to the method described by (33). LysM was stored at -80 °C, -20 °C and 4 °C for one year, or incubated at 25 °C and 30 °C for one month, and heated at 100 °C for 5 min; or kept in 50 mmol/L phosphate-citrate, Tris-HCl, and carbonic acid-bicarbonate buffers at pH values ranging from 2.0 to 12.0 for 24 h. The relative lytic activity of LysM after various treatments was evaluated using the above turbidity method, and then normalized to the level of fresh untreated LysM

### Crystal violet assay and calculation of biofilm disruption

Biofilm biomass quantified by a crystal violet (CV) microtiter plate assay was performed as previously described (27). Overnight cultures were diluted 1:100 into fresh medium and inoculated into wells of microtiter plates. For biofilm disruption, preformed biofilms were washed gently with PBS to remove planktonic cells and then exposed to the indicated treatment at 30 °C for the desired period. After treatment, the wells were stained with 0.1% crystal violet for 15 min at room temperature, rinsed three times with water. The bound dye was solubilized with 200 μL of 30% acetic acid for 10 min, transferred to a new flat-bottom plate, and absorbance was measured at 560 nm. Untreated biofilms served as controls. Biofilm disruption was calculated using the following equation:

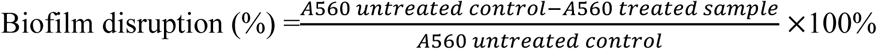

### Bacterial killing assay of LysM

The lytic activity of LysM and EDTA on planktonic bacteria was further determined by measuring the changes OD_600 a_nd bacterial CFU as described previously (33). Overnight bacterial culture of PAO1 was inoculated in fresh LB and incubated at 37 °C for 3 h, then collected and diluted in PBS to reach OD_600 o_f 0.6. 20 μL of 5 mM EDTA and/or 20 μL of 100 μg/mL LysM was added to 160 μL of bacterial culture to attain the desired concentration of EDTA (0.5 mM) and LysM (10 μg/mL), and the same volume of PBS was used as control, then incubated at 37 °C for 2.5 h. The OD_600 w_as examined every 30 min after treatment, and bacterial CFUs were measured before and after 2 h treatment by the plating method and colonies were counted after overnight incubation at 37 °C. *A. baumannii*, *K. pneumoniae*, *E. coli*, and *P. putida* were also supplied for this assay under the same conditions, and bacterial CFUs were measured before and after 2 h treatment. The bacterial killing rate (%) was calculated by using the following equation:

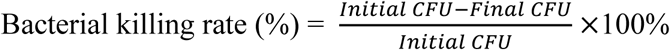

### Biofilm eradication and Biofilm bacteria killing assay of LysM

Biofilm-associated bacterial killing by LysM and EDTA was determined by colony-forming unit (CFU) enumeration as described previously with modifications (36, 37). Preformed biofilms of PAO1 were established in 96-well polystyrene plates in Jensen’s medium at 30 °C for 24 h, whereas A. baumannii, *K. pneumoniae*, *E. coli*, and *P. putida* biofilms were formed in LB at 30 °C under the same plate format. The wells with formed biofilm were gently washed three times with sterile PBS to remove planktonic cells. PBS containing EDTA (0.5 mM) and/or LysM was then added, and the plates were incubated at 30 °C for 2 h. Control wells received the same volume of PBS. CFUs were determined from parallel wells immediately before treatment and after 2 h of treatment. After treatment, the supernatant was removed and the remaining surface-associated biofilm was recovered in 1 mL of sterile saline by repeated pipetting and mechanical scraping, followed by vigorous vortex for 1 min. The recovered suspensions were serially 10-fold diluted, plated on LB agar plates, and incubated overnight at 37 °C for colony counting. The biofilm bacteria killing rate (%) was calculated using the following equation:

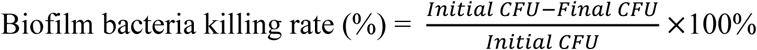

As an additional antibiofilm efficacy metric, killing can also be expressed as log reduction = log10(Initial CFU) − log10(Final CFU).

The efficacy of LysM and LysM-based cocktails to eradicate mature biofilms was evaluated as following. PAO1 biofilm was formed as described above by inoculating 1% of overnight bacterial culture in Jensen’s medium and incubating at 30 °C for 24 h without agitation in a polyvinyl chloride plate (costar). Following biofilm formation, the supernatants were removed, then each well was washed three times, and refilled with different solutions containing LysM (concentrations ranging from 6.25-50 μg/mL) and/or EDTA (0.5 mM) and/or PslG (50 nM) at 200 μL/well, PBS was used as control. After incubation at 37 °C for 2 h, supernatants were carefully removed, wells were washed three times, the remaining biofilm was measured by crystal violet staining as above. For live bacteria quantification post treatment, biofilm cells were recovered in PBS after various treatments by scraping the surface of the well with a pipette tip and rinsing with PBS. The recovered cell suspension was serially diluted with PBS, and viable biofilm counts were determined and calculated by plating method as described above.

### Determination of minimum inhibitory concentration (MIC) and minimal biofilm-inhibition concentration (MBIC)

The inhibitory effects of LysM on the growth of *P. aeruginosa* were evaluated by measuring MIC and MBIC, which represent the minimum concentration of antibiotic required to inhibit the growth of free bacteria and biofilm formation, using microtiter test as described with slight changes (38). Briefly, the overnight bacterial culture of PAO1 was inoculated in Jensen’s medium at a concentration of 1%, then the bacterial solution was added to a 96-well polyvinyl chloride plates (costar) at 100 μL/well. Double serial dilutions of the LysM in Jensen’s medium were applied in a concentration range of 100-0 μg/mL for testing wells, and incubated at 30 °C for 24 h without agitation. The planktonic culture was carefully moved to a new microtiter plate subjected for the measurement of OD_600 t_o determine MIC value. The remaining biofilm formed on each well was gently washed three times, followed by staining with 0.1% crystal violet for 30 min at room temperature. The adherent stain was solubilized in 30% acetic acid and OD_560 nm w_as measured to quantify the biofilm biomass.

### Flow-cell biofilm cultivation and treatment

As described previously (39), *P. aeruginosa* biofilms were grown in Jensen’s medium at room temperature in three-channel flow chambers (1×4×40 mm). Mid-log-phase cultures (OD600 = 0.5; approximately 10^7^ cells) grown in Jensen’s medium were used for inoculation. After inoculation, the medium flow was stopped for 1 h to allow initial attachment and then resumed at 3 mL/h of Jensen’s medium flow with or without LysM. The biofilms were further cultured under continuous-flow conditions for 12 h. Images were acquired by Nikon Ti2 with 3 Second interval and 20000 frames were obtained. Matlab 2022b was used for data analysis with program biotree. The bacteria number in per frame were counted, denoted as *N_i_*, and the cumulative bacterial number (N) in all frames were calculated using the following equation:

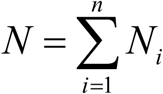

The trajectory and the fate of bacteria were obtained by biotree.

### Biofilm cultivated in microfluidic device and image acquisition

The biofilm in microfluidic device was cultivated as described by Jintao Liu (32). To grow a *P. aeruginosa* biofilm, an overnight culture of PAO1 was used for inoculation and Jensen’s medium was utilized to grow biofilm. LysM solution 500 μL (3 mg/mL) or PBS buffer was added to 50 mL of medium after approximately 100 bacterial cells being trapped at the designed seeding zone in a microfluidic device. The flow rate was approximately 500 μm/s, and the temperature was kept at 37 °C. To cultivate a *E. coli* biofilm in the microfluidic device, M63B1 medium (100 mM KH_2P_O_4,_ 15 mM (NH_4)2S_O_4,_ 0.8 mM MgSO_4,_ 3 μM vitamin B1, 22 mM glucose, adjusted to pH 7.4 with KOH) was used. PI was suspended in the culture medium at 1:1000 dilution to stain dead bacteria. After 20 h of growth under the medium flow, LysM and EDTA was added to the loading medium to final concentrations of 30 μg/mL and 0.5 mM, respectively in order to examine their effect on a preformed *E. coli* biofilm. The same volume of PBS was added to un-treated biofilm, then incubated at 37 °C for another 20 h. The growth of biofilm was tracked by time-lapse images acquired via an Olympus IX83 microscope (Andor Zyla 4.2 sCMOS camera) and analyzed using ImageJ and MATLAB to quantify biofilm growth.

### Pellicles cultivation and image acquisition

Pellicles (Air-liquid interface biofilms) were grown in glass chambers (Chambered #1.5 German Coverglass System, Nunc) as described previously (27). One milliliter of a 1% dilution of PAO1 overnight culture in Jensen’s medium was transferred into a well of the chamber, incubated at 30 °C for 24 h. The media were soaked out and replaced by various treatments of EDTA, EDTA+LysM, PslG+EDTA+LysM, PslG or PBS. The working concentration of EDTA, LysM, and PslG was 0.5 mM, 10 μg/mL, and 50 nM respectively. Then the corresponding treating solution was removed post 2 h-treatment, and the remaining biofilms were gently washed three times with PBS, and were stained with SYTO9 (5 µM, molecular PROBE, Invitrogen) and propidium iodide (PI, 10 µM, YEASEN) for 15 min, followed by washing before observation. The images were acquired using an FV1000 Confocal laser scanning microscopy (CLSM, Olympus, Japan) with the 63×/1.3 objective at Ex/Em of 483/503 nm (SYTO9) and 493/636 nm (PI), and then processed using COMSTAT software.

### Fluorescence and Phase contrast microscopy

Cultures for single-cell analysis were prepared as described above, the exponential phase bacterial suspension (PI was added) with or without LysM and EDTA was grown in an incubator at 37 °C. The 30 μg/mL LysM and 0.5 mM EDTA was used as the working concentration, respectively. After one-hour inoculation, the samples were ultrasonicated briefly to disperse aggregated cells, 5 μL of the culture was withdrawn and placed on a glass coverslip, then phase and fluorescence images were collected as following. Phase-contrast and microscopy images of the slide were collected on a Nikon TE2000U inverted fluorescence microscope equipped with a cryocooled Coolsnap HQ2 charge-coupled-device camera and a GFP or red fluorescent protein (for PI) filter set. The fluorescence and phase-contrast microscopy images were analyzed by using a previously described method (40, 41).

### A549 lung epithelial cells cytotoxicity assay

Cell cytotoxicity of LysM was performed on A549 lung epithelial cells by cell counting kit-8 (CCK-8, Dojindo, Japan) as described previously (23). Briefly, A549 lung epithelial cells were seeded to a 24-well plate at 2.5×10^5^ cells/well and incubated at 37 °C, 5% CO_2,_ for 12 h, followed by incubation in fresh DMEM-FAS medium containing various concentrations of LysM for another 12 h. Then, cells were stained with CCK-8 and incubated according to instruction. The optical density at 450 nm was measured using Synergy H1 Hybrid microplate reader (BioTek Instruments, Inc.). The assays were conducted three times independently.

### HAOs culture

Human nasal epithelial biopsy samples were collected from patients admitted to the hospital. Written informed consent was obtained from all patients, and the study was approved by the Ethics Committee of the Seventh Affiliated Hospital of Sun Yat-sen University (approval number: KY-2021-075-02). As previously described (42, 43), Nasal biopsy samples were washed in PBS and digested for 5-10 min at 37 °C with a mixture of neutral protease, collagenase D, DNase I, and Accutase. After centrifugation, the cell suspension was passed through a cell strainer and washed with PBS at least three times. Then cells were cultured and passaged in a T75 flask at 37 °C, 5% CO_2,_ and 95% relative humidity using Pneumacult-Ex Plus medium (Stemcell Technologies, Canada, catalog number 05040). Once the cells reached 60-80% confluence, they were detached from the bottom of the culture flask using accutase dissociation solution (Sigma-Aldrich, Germany, catalog number A6964) and transferred to cell culture inserts with a permeable surface (0.33 cm^2^, 0.4-μm pore size; Corning, USA, catalog number 3470), seeding 5×10^4^cells per insert. The cells were cultured with Pneumacult-Ex Plus medium for 3 days until complete confluence was achieved. Pneumacult ALI medium (Stemcell Technologies, Canada, catalog number 05001) was utilized for cell growth and differentiation. The medium was refreshed every 48 h. After 18 days of differentiation, Human Airway Organoids (HAOs) were obtained and identified using H&E staining or immunofluorescence for subsequent use.

### Hematoxylin-eosin (H&E) staining

The HAOs were fixed overnight in aqueous 4% paraformaldehyde (PFA, Beyotime, PRC), washed with PBS, embedded in paraffin, sectioned at 3 µm, and mounted on slides. H&E staining was performed following a standard protocol as the previous study (43).

### TEER measurements

The Millicell ERS system (Millipore) is utilized to assess the transepithelial electrical resistance (TEER) of HAOs under normal and infected conditions. Before taking readings, HAOs were washed for free of mucus and placed in pre-warmed Pneumacult-ALI medium (200 µL apical / 400 µL basolateral). Following subtraction of the background signal from empty inserts, TEER (Ω·cm²) was calculated as:

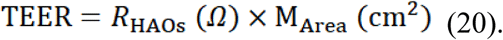

### Immunofluorescence

Immunofluorescence of paraffin-embedded HAO sections (3 µm thick) were conducted as previously described (42). After deparaffinization and rehydration, antigen retrieval was performed by heating the sections in a water bath with citrate buffer (10 mM; Beyotime, China) at 95 °C for 15 min. Subsequently, the HAO sections were washed three times in blocking solution (Solarbio, China). The sections were incubated with the primary antibody (INC) overnight at 4 °C. Afterward, the sections were washed three times in PBS containing 0.05% Tween 20 (PBST; Sangon Biotech, China), followed by incubation with the secondary antibody (INC) for 1 hour at room temperature. After three washes and counterstaining with 4,6-diamidino-2-phenylindole (DAPI; Solarbio, China), the samples were mounted using ProLong Gold Antifade Mountant (Invitrogen, USA). The samples were imaged using a CLSM (Leica TCS SP8, Germany). Primary antibodies were employed at a dilution of 1:200, including acetyl-α-tubulin (Lys40) (D20G3) Rabbit MAb (Cell Signaling Technology, 5335S) and MUC5AC (E3O9I) Mouse MAb (Abcam, ab3649). Secondary antibodies were used at a dilution of 1:500, including Goat anti-Rabbit IgG (H+L) Cross-Adsorbed Secondary Antibody, Alexa Fluor 488 (Invitrogen, A-11008), and Goat anti-Mouse IgG (H+L) Highly Cross-Adsorbed Secondary Antibody, Alexa Fluor 594 (Invitrogen, A-11032).

### Cultivation of *P. aeruginosa* biofilm on HAOs and evaluation of biofilm inhibition and disruption by LysM

*P. aeruginosa* PAO1::*mScarlet-I* strain was used for the HAOs infection. To prepare bacterial suspension to load on HAOs, an overnight PAO1 culture was diluted into fresh LB medium (1/100 dilution) and incubated at 37 °C with shaking at 200 rpm for 3 h, then bacteria were pelleted and suspended in PBS buffer. HAOs were gently washed three times with PBS (200 μL per wash) prior to bacterial inoculation. For the biofilm inhibition assay, 10 μL of bacterial suspension (1×10^5^ CFU) was first added onto each HAO, followed by 10 μL of various treatment solutions. The treatment groups included PBS alone, 30 μg/mL LysM in PBS, LysM+PslG (PBS containing 10 μg/mL LysM and 50 nM PslG), LysM+EDTA+PslG (PBS containing 10 μg/mL LysM, 0.5 mM EDTA, and 50 nM PslG), and PslG+CIP (PBS containing 50 nM PslG and 1 μg/mL CIP). The infected HAOs were incubated at 37 °C in 5% CO_2 f_or 24 h. Biofilms on HAOs were washed gently for three times with PBS and imaged with CLSM (Leica TCS SP8 microscope, Germany). Following CLSM imaging, HAOs were examined by H&E staining or immunofluorescence. For the biofilm disruption assay, HAOs were first infected with 20 μL of bacterial suspension containing 1×10^5^ CFU of PAO1::*mScarlet-I* and incubated at 37 °C in 5% CO_2 f_or 24 h to allow biofilm formation. Next, the same treatments described above were applied and treated for 6 h at 37 °C in 5% CO_2._ After treatment, the organoids were washed three times with PBS and biofilm were analyzed by CLSM.

### Checkerboard experiment-synergy of antibiotics and LysM

To evaluate the synergistic effects of LysM and antibiotics, chequerboard assay was conducted by using the 96-well-plates method as described (44) with slight modifications, and tetracycline (TET), erythromycin (ERY), carbenicillin (CRB), colistin, ciprofloxacin (CIP), tobramycin (TOB) were applied for this assay. Briefly, the overnight bacterial culture of PAO1 was diluted to OD_600 o_f 0.1, then 1% inoculated into fresh LB containing 2-fold dilutions of each antibiotic or/and LysM in a 6×8 dose-point matrix. The plates were incubated at 37 °C overnight, and the absorbance of OD_600_ was measured. At least three biological replicates were completed for each combination. Heat maps were generated according to OD_600,_ and the FIC indices (FICi) for LysM with each antibiotic were calculated. The interactions were interpreted based on the calculation of FICi, in which: FICi ≤ 0.5, synergistic; 0.5 < FICi ≤ 1, additive; 1< FICi ≤ 2, neutral; and FICi ⩽ 2, antagonistic, respectively.

### Anti-biofilm effect of LysM-based cocktail against clinical *P. aeruginosa* isolates

The clinical *P. aeruginosa* isolates M3, M28, CC2, CC3, CC4, CC7, CC11, and CC12 were selected to evaluate the antibiofilm efficacy of LysM cocktail (PslG+LysM+EDTA) against MDR *P. aeruginosa*. The biofilm biomass and biofilm bacteria were measured as described above, concentrations of 50 nM, 10 μg/mL and 0.5 mM were used as the working concentration of PslG, LysM, and EDTA, respectively, and the bacteria killing rate was calculated as described above.

### *G. mellonella* infection model

Healthy *G. mellonella* larvae (weighing 250–350 mg) were randomly divided into groups (n=10 per group) and infected with 10 μL of bacterial suspension (5×10^2^ CFU/larva) at the second right posterior proleg. At 20 min post-infection, larvae were treated with LysM alone or combined with different antibiotics. Survival rates of *G. mellonella* were recorded every 2 h for 28 h. Survival curves were plotted using the Kaplan–Meier method and compared using the log-rank test.

### Statistical analysis

All the data except of *G. mellonella* survival rates were analyzed by using GraphPad Prism 8, and Student’s *t*-test was used to determine statistical significance of *P*-values. \*\*\**P* < 0.001, \*\**P* < 0.01, \**P* < 0.05. Error bars shown in the pictures represent ± SD (standard deviation) from at least three biological replicates. The statistical analysis of *G. mellonella* survival rates were determined by the two-sided, log [rank] (Mantel-Cox) test.

## Acknowledgement

We thank Dr. Wenbin Du at Institute of Microbiology, Chinese Academy of Sciences for the support on microscopy.

## Video Legends

**Video 1.** An example video of the morphological transition and lysis of *P. aeruginosa* PAO1 cells during LysM and EDTA treatment. The video was taken at a frame interval of 30 s for 30 min and was played back at 5 fps.

**Video 2.** An example video of tracking the biofilm formation of *P. aeruginosa* PAO1 in a flow-cell system without (A) or with LysM treatment (B) during the first 2 h of treatment. The video was taken at a frame interval of 1 min for 2 h and was played back at 5 fps.

**Video 3.** An example video of tracking the expansion of biofilm formed by *P. aeruginosa* PAO1 in a microfluidic device under a Jensen’s media flow without (A) or with LysM (B). The video was taken at a frame interval of 5 min for 24 h and was played back at 5 fps.

**Video 4.** An example video of tracking the expansion of *E. coli* biofilm formed in a microfluidic device. **A**. Biofilm grown under media flow; **B.** Biofilm grown under media flow with LysM and EDTA, which was added at 20 h post inoculation. The video was taken at a frame interval of 5 min for 40 h and was played back at 5 fps. Shown was the video from 21 h to 40 h post inoculation.

## Extended Data

### Legends of supplemental figures

**Fig. S1 The addition of LysM leads to an explosive cell lysis with the help of EDTA. A.** Morphology changes of *P. aeruginosa* PAO1 after the treatments for 30 min: no treatment (PBS), LysM, LysM+EDTA. The photographs were taken at 400 × magnification using a phase-contrast microscope with crystal violet (CV) staining after treatments. **B.** Visualization of explosive cell death of *P. aeruginosa* PAO1 by the time-lapse microscopy, and LysM with EDTA was added at the beginning. Yellow arrows represent the process of explosive cell death. **C-D.** Effects of LysM with EDTA on the cell death of *P. aeruginosa* PAO1. Microscopy images of bacteria post 30 min of treatment (C), images of brightfield and red fluorescence (PI-stained dead cell) were shown. Histograms of red fluorescent signal of individual bacterial cell measured after 30 min of treatment. Fluorescence per cell was measured as described in (41). The length of each horizontal bar indicates the percentage of cells that fluoresced at the indicated level. The heavy horizontal line in the scatter plot indicates the median fluorescence of the population. Scatter plot represents red fluorescence of individual bacterial cells with or without treatment.

**Fig. S2 Stability of LysM and 3D structural superposition of its mutations. A.** Stability of LysM under different pH from 2 to 12. **B.** Storage stability of LysM under various temperatures (-80 °C, -20 °C and 4 °C for one year, 25 °C and 30 °C for one month, 100 °C for 5 min). Error bars indicate the mean±SD of three independent experiments. Statistical significance was determined using Student’s *t-*test (*** *P* < 0.001, * *P* < 0.05, ns, non-significant). **C.** 3D structural superposition of the wild-type LysM (PA0629, light blue) and its point mutants E51A (purple), E60A (orange), L126A (salmon), W165A (wheat), and R186A (yellow). The overlaid ribbons show that the mutations do not significantly affect the overall three-dimensional folding of the protein.

**Fig. S3 LysM homologues analysis. A**. Heat map showed the source of LysM homologues with identity of amino acid sequence over 60%. **B**. Amino acid sequence alignment of LysM with its homologues found in bacterial and phage genome. The alignment was carried out by T-coffee, and the results were generated using ESPript. Residues conserved in all sequences were highlighted in red.

**Fig. S4 The effect of LysM and its “cocktail” on bacterial growth and biofilms of *P. aeruginosa* and *E. coli*. A.** The living bacteria in *P. aeruginosa* PAO1 biofilm were measured after 2-h or 6-h treatment of the LysM “cocktail”, TOB and CIP of MIC or 2×MIC combined with PslG were used as controls (MIC of TOB and CIP is 2 and 1 μg/mL, respectively). **B.** The corresponding living bacteria in biofilm under treatment of LysM at 6.25 μg/mL (MBIC) with/without EDTA (0.5 mM) shown in Fig. 4A. **C.** The corresponding CFU of planktonic bacterial under treatment of LysM at 6.25 μg/mL with/without EDTA (0.5 mM) shown in Fig. 4B. **D.** Biofilm biomass and the living bacteria in biofilm of *E. coli* after treatments in microtiter dish. The 20 h-old biofilm of *E. coli* was exposed to LB supplemented with EDTA and/or LysM (30 μg/mL) at 37 °C for 2 h and then biofilm biomass and CFU were measured. Error bars indicate the mean ±SD of three independent experiments. Statistical significance was determined using Student’s *t-*test (*** *P* < 0.001, ns, non-significant).

**Fig. S5 LysM inhibits the bacterial growth and biofilm formation of *A. baumannii*, *E. cloacae*, *K. pneumoniae*, *E. coli*, and *P. putida*.** The biofilms of *A. baumannii*, *E. cloacae*, *K. pneumoniae*, *E. coli* and *P. putida* were treated with PBS containing LysM and/or EDTA for 2 h. The biofilm disruption rate (%) was shown in A and the corresponding CFU of living bacteria in biofilm post treatment was shown in B. Error bars indicate the mean±SD of three independent experiments. Error bars indicate the mean±SD of three independent experiments. Statistical significance was determined using Student’s *t-*test (*** *P* < 0.001, ns, non-significant).

**Fig. S6 The cytotoxicity of LysM, EDTA, PslG and LysM+EDTA+PslG “cocktail” at different concentrations. A-D.** The cytotoxicity of LysM (A), EDTA (B), PslG (C), LysM cocktail (D) at different concentrations. The cell viability of A549 lung epithelial cell detected by the CCK-8 assay. Error bars indicate the mean ± SD of three independent experiments. Statistical significance was determined using Student’s *t-*test (ns, non-significant).

**Fig. S7 LysM “cocktail” inhibits biofilm formation and disrupts the mature biofilm of *P. aeruginosa* on HAOs.** Visualization of biofilm architecture, tissue/cellular morphology, and spatial distribution of human airway organoids (HAOs) under different treatment groups, namely PBS, EDTA, LysM+EDTA, PslG, and PslG+EDTA. For each treatment group, biofilms were visualized by CLSM (left panels). HAO morphology was assessed by H&E (middle panels). The expression and localization of epithelial markers acetyl-α-tubulin (cilia) and MUC5AC (mucin) were detected by immunofluorescence, with nuclei counterstained by DAPI (blue). Scale bars = 10 μm.

**Fig. S8. Dose-dependent *in vivo* protection and broad-spectrum re-sensitization of MDR clinical isolates of *P. aeruginosa* by LysM**. **A.** LysM restores the antibiotic susceptibility of MDR clinical isolates of *P. aeruginosa* (CC7 and CC11). Left and Middle: Growth curves demonstrating the potentiating effect of LysM on the MIC of CIP against CC7 and CC11, respectively. Right: Potentiating effect of LysM on the MIC of Colistin against CC11. The solid arrows indicate the shift in MIC values upon the addition of LysM. **B.** Survival curves of *G. mellonella* larvae infected with *P. aeruginosa* PAO1. Left: Dose-dependent protective efficacy of LysM monotherapy at 1, 2, and 4 μg/mL. Middle: Synergistic *in vivo* therapeutic effect of LysM (1 μg/mL) with the PslG (50 nM). Right: Comparison of the therapeutic efficacy between the antibiotic-free, dual-enzyme combination (LysM+PslG) and various LysM+antibiotic regimens.

### Legends of supplemental Tables

**Table S1 List of the information of all strains, plasmids, primers, chemicals and detection kits used in this study.**

**Table S2 The resistant pattern of durg-resistant *P. aeruginosa* isolates**

